# High-throughput High Content Quantification of HIV-1 Viral Infectious Output

**DOI:** 10.1101/2025.06.28.662157

**Authors:** Teresa LuPone, Alexis Brantly, Oluwatofunmi Oteju, Stephanie M. Matt, Kaitlyn Runner, Emily Nickoloff-Bybel, Micheal Nonnemacher, Peter J. Gaskill

## Abstract

Infection with human immunodeficiency virus (HIV-1) remains a global health issue and still drives the development of significant pathology and various comorbidities. Antiretroviral therapy (ART) can effectively suppress viral replication but is often initiated months or years after initial infection, leaving a substantial period in which viral replication progresses unchecked. While ART suppresses HIV-1 replication, it does not prohibit the development of HIV-1-associated comorbidities, highlighting a lack of understanding in the connection between replication and HIV-1-associated pathogeneses. Thus, it is critical to better define HIV-1 replication dynamics to more effectively target different stages of the viral replication cycle in distinct cell populations. Here, we show a high-content imaging reporter assay that uses modified human osteosarcoma cells expressing HIV-1 receptors (GHOST cells) which fluoresce in response to HIV-1 infection. These cells have been previously used to assess HIV-1 infectivity and tropism, but this modified assay enables rapid evaluation of large numbers of samples with consistency and replicability, while also easily integrating into existing experimental pipelines that analyze p24 secretion in collected supernatants. This also allows for direct correlation between infectivity and p24 secretion, resulting in a deeper interrogation and more robust understanding of HIV-1 infection kinetics.

**Institutional Permissions:** All of the primary cells used in this study were obtained from donors in accordance with Institutional Review Board protocols at the New York Blood Center and Drexel University. All studies using primary cells were performed with cells from deidentified donors and were found by the Institutional Review Board at Drexel University to be exempt from human subjects protocols (protocol number 2208009386). All experiments using *in vitro* cell lines or primary human cells were approved by the Institutional Biosafety Committee and at Drexel University.

**Highlights:**

- The current toolkit for evaluating *in vitro* HIV-1 infection dynamics does not currently include an assay providing direct, efficient assessments of infectivity in large numbers of samples.
- Direct assessment of viral infectivity longitudinally alongside other measurements of viral infection such as p24 secretion or amount of proviral DNA enables correlation of different stages of viral replication efficiency over time.
- The high-content viral titer assay described here correlates with widely used surrogate measures of viral infectivity, such as p24 secretion, and integrates easily into existing experimental pipelines that collect supernatant from infected cultures.
- Adapting fluorescent reporter assays to a high-content imaging and analysis pipeline creates a high-throughput assay of direct infectivity that enables evaluation of changes in infectivity across multiple treatments or timepoints and integrates easily into existing experimental pipelines that collect supernatant from infected cultures.

## 1. Introduction

Globally, more than 39 million people are living with human immunodeficiency virus (PWH)^1^, of which the majority have access to anti-retroviral therapy (ART). ART greatly decreases viral replication, but the initiation of this therapy is often several years after infection^2^ and ART does not eliminate virus from infected cells. Irrespective of the viral secretion, HIV-1 infected cells can continue to produce inflammatory and viral factors that contribute to chronic inflammation^3–5^ and tissue specific pathologies^6–10^, driving the comorbidities now commonly associated with chronic HIV-1 infection. The consistent presence of HIV-1 in anatomical reservoirs, particularly myeloid populations in the brain, lungs, liver and gut, likely also contributes to these long-term effects. To better identify and define the kinetics and spread of infection within these populations, it is critical to understand HIV-1 replication kinetics with these cell types.

Traditionally, HIV-1 replication has been evaluated *in vitro* using several types of assays; measurements of viral entry (appearance of viral marker within a target cell), the production of viral RNA (viral transcription), the production and secretion of viral proteins (viral budding, protein secretion), and the secretion and/or maturation of infectious virions (viral titer, reverse transcriptase activity) (Figure 1). These assays can also be classified by how they measure viral output; either indirectly, measuring surrogates of infection, or directly, measuring infectivity in target cells. Currently, the most common assays for evaluating HIV-1 replication are indirect, using PCR assessment of viral RNA or measuring secretion of the viral capsid protein (p24Gag) via an ELISA or similar assay ^11–18^. Evaluation of reverse transcriptase (RT) activity in secreted virions is also used, though predominantly in T-cell cultures ^19–27^.

**Figure 1.**
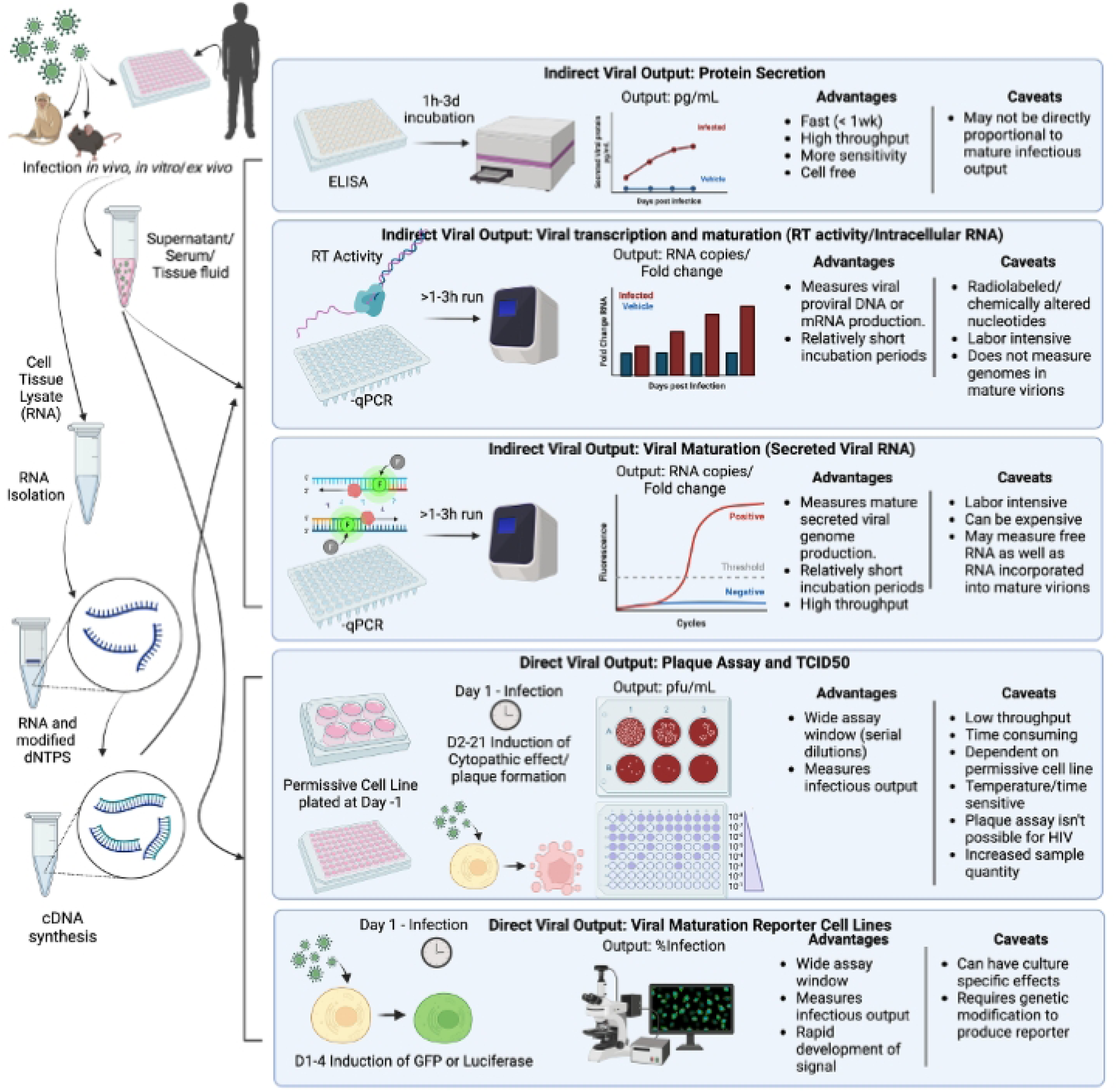
Classical methods used to calculate viral output contain advantages and disadvantages in evaluation of infectivity. In general, experiments are performed in vivo, ex vivo, and/or in vitro with HIV-1 and samples are collected on different days post infection including supernatants, protein, and RNA lysates. Indirect viral quantification methods are broken down into 3 categories: protein secretion, viral transcription, and viral maturation. Indirect viral quantification methods include ELISA-based protein secretion detection, RT activity, and qPCR measurements of viral gene expression. Direct viral quantification methods include: plaque assays and TCID_50_, as well as infectivity assays using permissive cell lines. Created with BioRender.com.

Direct assessments of viral output are traditionally end-point assays that use a visual cytopathic effect to evaluate the number of mature infectious virions. Plaque assays are among the most common of these, generally adding virus to a plate of cells and enumerating the number of plaques – areas of dead cells – produced when the virus forms syncytia or kills infected cells via lytic infection^17, 28–30^. The readout for these is plaque forming units (PFU). Other direct infectivity assays capitalize on effects such as cell fusion or lytic cell death and are measured by the amount of virus that kills 50% of the target cell population (TCID_50_)^31–37^. Plaque assays are ineffective for HIV-1, and TCID_50_ assays for HIV-1 can be quite labor intensive, as HIV-1 inoculum must be serially diluted to acquire an effective assay window. Additionally, production often varies between cultures of the same cell line and even more so in primary cells from different donors, necessitating longer-term culture of many wells and/or plates per sample. This makes it time– and resource intensive to assay large numbers of samples or conditions.

To accelerate these assays, reporter cell lines that produce GFP or luciferase signal upon HIV-1 infection been developed^38, 39^. These include luminescent TZM-bL cells^40^ and fluorescent lines such as JLTRG-R5, GHOST, and CEM-GFP. Cells for these assays are often genetically modified to express HIV-1 receptors and/or co-receptors to allow or increase infectability, or to measure specific tropism^39, 41, 42^. This study used GHOST cells, human osteosarcoma cells stably expressing CD4 and either CCR5, CXCR4, and/or other co-receptors^43–45^. Originally developed to assess HIV-1 co-receptor tropism^44^, GHOST cells also contain a Tat-driven HIV-1-2 LTR fused with GFP and fluoresce at 509 nm within 48 hours post infection with HIV-1^46^. Reporter lines like these can accelerate evaluation of direct infectivity but still require the assessment of a large number of infected cultures to establish a clear assay window and introduce potential experimenter bias when using manual microscopy.

In this study, we addressed these concerns by optimizing the analysis of the GHOST reporter line on a CellInsight CX7 high-content screening platform, enabling the accurate, unbiased acquisition and analysis of an entire 96 well plate of infected GHOST cells within 1 to 2 hours^47–49^. Traditional viral titer assays using GHOST cells in 6-well plates were first adapted to 96 well plates, enabling analysis of more samples using a smaller number of plates in less time, while also reducing experimenter bias by automating the acquisition and image analysis. We then demonstrated the utility of this assay in assessing the infectivity of HIV-1 viral stocks derived from commonly used lab-adapted strains and participant derived viruses, as well as the assessment of longitudinal changes in the infectivity of secreted virions released from infected cultures of primary human monocyte-derived macrophages.

## 2. Methods

### 2.1 Reagents

Cell culture reagents including RPMI-1640 and DMEM media, non-essential amino acids, penicillin/streptomycin (P/S) and Trypsin 0.05% + EDTA were obtained from Invitrogen (ThermoFisher, Carlsbad, CA, USA). Bovine serum albumin (BSA) and glycine from Sigma-Aldrich (cat # A7906-100G, St. Louis, MO, USA). Tween, dimethyl sulfoxide (DMSO), and hydroxyethyl piperazineethanesulfonic acid (HEPES) were obtained from Fisher Scientific (Hampton, NH, USA). Geneticin (G418) and Hygromycin were obtained from ThermoFisher. Puromycin was obtained from R&D Systems (Minneapolis, MN, USA). Fetal bovine serum (FBS) was from Fisher Scientific and human AB serum was from Gemini Bio-Products (Invitrogen, ThermoFisher,). Tris Buffered Saline (TBS) Tween 20 was obtained from Fisher Scientific. Macrophage colony stimulating factor (M-CSF), was from Peprotech (Rocky Hill, NJ, USA).

### 2.2 Cell lines

GHOST (3) Parental (ARP-3679) and CCR5+ Cells (Hi-5, ARP-3944), were obtained through the NIH HIV Reagent Program, Division of AIDS, NIAID, NIH: GHOST (3) Parental, ARP-3697, and CCR3+ CXCR4+ CCR5+ Cells, ARP-3944, contributed by Dr. Vineet N. KewalRamani and Dr. Dan R. Littman. Parental Ghost cells are cultured in complete DMEM containing, 10% FBS, 500μg/ml G418, 100μg/ml Hygromycin, 1% P/S. Ghost Hi5 cells are cultured in the same media as parental cells with the addition of 1μg/ml Puromycin as a selection marker. CEM•SS cells (ARP-776) were obtained through the NIH HIV-1 Reagent Program, Division of AIDS, NIAID, NIH: CEM•SS Cells, contributed by Dr. Peter L. Nara. CEM•SS cells were cultured in RPMI 1640 10% FBS, 1%P/S and 1% HEPES. Lenti-X cells (Takara Bio USA Inc, Takara Bio Inc, Shiga, Japan) were cultured in DMEM containing 10% FBS, 1% P/S, and 1% Nonessential Amino acids.

### 2.3 Isolation, culture, and infection of primary human monocyte derived macrophages (hMDM)

Human peripheral blood mononuclear cells (PBMC) were separated from blood obtained from de-identified healthy donors (either the New York Blood Center (NYBC), Long Island City, NY, USA or the Comprehensive NeuroHIV-1 Center (CNHC) and Clinical and Translational Research Support Core (CNHC/CTRSC) Cohort at Drexel University College of Medicine, Philadelphia, PA, USA) by Ficoll-Paque (GE Healthcare, Piscataway, NJ, USA) gradient centrifugation. Blood collection was conducted in accordance with the protocols approved by the Institutional Review Board (IRB) at the NYBC. Blood collection by the CNHC/CTRSC was approved by the IRB at Drexel University, IRB protocol 1609004807. All experiments with material isolated from human blood were performed using protocols approved by the Institutional Biosafety Committee at Drexel University.

Following isolation of PBMC, monocyte percentage was determined using a Pan Monocyte Isolation Kit (Miltenyi Biotec, Auburn, CA, USA) and PBMC were plated in 96 well plates at 15,000 monocytes per well. Cells were cultured in macrophage media (DMEM + Glutamax with 10% FBS, 5% human AB serum, 10 mM HEPES, 1% P/S, and 10 ng/mL M-CSF), replacing media with fresh media at day 3 post-isolation. Cells were considered mature after 6 days in culture, and all infections were performed on days 6 and 7 post-isolation. Mature hMDM were inoculated with 1 ng/mL of HIV-1_ADA_, HIV-1_YU2A_ or HIV-1_YU2B_, or patient-derived strains of HIV-1 (9201R1, THRO, CH167) for 24 hours, then washed to remove excess virus and given fresh macrophage media. The hMDMs remained in culture for 9 days after initial inoculation, collecting supernatant at 3, 5, 7 and 9 days post infection (DPI). Supernatants from all wells in a single experimental condition were collected, pooled and aliquoted for storage at –80°C. Concentration of p24Gag protein (p24) in supernatant is determined by HIV-1 p24 high sensitivity AlphaLISA Detection kit (AL291C, Revvity, Waltham, MA) and infectivity of secreted virions was determined using the viral titer assay described below.

### 2.4 Generation of Infectious HIV-1 viral stocks

Viral stocks were generated by infecting CEM-SS cells with a blood derived, R5-tropic strain of HIV-1 (HIV-1 ^50^), as we have done previously ^51, 52^. Briefly, CEM-SS cells cultured in T175 flasks in RPMI-1640 containing 10% FBS and 1% P/S. Culture was inoculated with a single tube of HIV-1_ADA_ obtained through the NIH HIV-1 Reagent Program, Division of AIDS, NIAID, NIH: Human Immunodeficiency Virus-1 ADA, ARP-416, contributed by Dr. Howard Gendelman. Infected cultures were maintained until syncytia formation was observed, generally after 15 – 18 days. Once syncytia were observed, supernatant was collected every 24hr until viral replication outpaced cell growth, killing the culture. The stocks used in this study showed syncytia at 17 days post-infection, and cell-free supernatants were collected daily from 18 to 41 days post-infection. Supernatants were spun down to remove cells and debris, then transferred to a fresh conical vial, aliquoted in 1 mL aliquots, and stored at –80°C for use as viral stocks. The concentration of viral stock was determined by quantifying the amount of p24•Gag per mL, using the HIV-1 p24 high sensitivity AlphaLISA Detection kit (Revvity).

Viral stocks of all HIV-1 strains other than HIV-1_ADA_ were generated by transfection of the Lenti-X 293T cell line (Takara Bio USA Inc, Takara Bio Inc, Shiga, Japan). Lenti-X cells are a HEK293T line transformed with adenovirus type 5 DNA and expressing SV40 large T antigen that is optimized for lentiviral vector production. The plasmid containing the R5-tropic strain HIV-1_YU2**A**_ (ARP-1350) was obtained through the NIH HIV-1 Reagent Program, Division of AIDS, NIAID, NIH: Human Immunodeficiency Virus 1 (HIV-1) YU2 Infectious Molecular Clone, contributed by Dr. Beatrice Hahn and Dr. George M. Shaw. Plasmids containing full-length R5-tropic strains HIV-1_CH167_, HIV-1_9201R1_, HIV-1_THRO_, HIV-1_YU2**B**_ were a generous gift from Dr. Katharine J. Bar at the University of Pennsylvania. Importantly, Figure 5 uses HIV-1_YU2_ obtained from two distinct sources – the NIH HIV-1 Reagent program and Katherine J. Bar – these are labeled HIV-1_YU2**A**_ and HIV-1_YU2**B**_, respectively. The HIV-1_YU2**A**_ plasmid was amplified by Genescript (Piscataway, NJ, USA), while the HIV-1_YU2**B**_ plasmid, as well as all the plasmids containing participant-derived viruses, were amplified by the Bar lab. Following amplification, all plasmids were diluted to a concentration of 1mg/mL in H_2_O, aliquoted and stored at –20°C.

Lenti-X cells cultured as described above were grown until 70% confluent in T175 flasks, then plated on 10cm^2^ dishes (Fisher-Scientific) at 2 x 10^6^cells/plate and maintained for 2 to 3 days until plates reached 80-90% confluency. Viral transfectants were prepared by adding 3 μg viral of plasmid (3μL at 1 mg/mL) to 36 μL of FugeneHD (Roche, Basel, Switzerland) and 800 μL of Opti-MEM (ThermoFisher), vortexing for 5 – 10 seconds, and incubating at room temperature for 25 min. During incubation, Lenti-X plates were trypsinized using 2mL 0.05% Trypsin EDTA (Invitrogen, ThermoFisher) at 37°C for 5 min, then cells were gently lifted and moved into a 15mL conical, adding 6 mL fresh culture media to neutralize the trypsin. Cells were then counted using a Nexcelom Cellometer Auto 2000 Cell Viability Counter (Nexcelom Biosciences, Waltham, MA, USA) using equal volumes of cells and AOPI (Nexcelom Biosciences), centrifuged at 1000 rpm for 5 min at RT and resuspended at 1×10^6^cells/mL.

After a 25 min incubation, the Opti-MEM-FugeneHD-plasmid viral transfectant mix is dotted onto fresh 10cm^2^ dishes, and Lenti-X cells are added dropwise on top of the mixture to maximize surface area contact between cells and plasmid mix. Cells are plated to a final density of 8-10 x 10^6^ cells/plate in 1 mL, then 9 mL of Lenti-X media is added to each plate. Each transfection was incubated for 24 hrs at 37°C, washed and given fresh media. Plates were then incubated for 72 hrs and supernatants from all plates are pooled in a single 50 mL conical vial, and incubated with a PEG-IT 5x solution (System BioSciences, Palo Alto, CA, USA) on an orbital rocker at 4°C for 24 hrs. After incubation, pegylated virus was pelleted by centrifugation for 30 min at 1500 x g at 4°C. Following centrifugation, media was aspirated and pegylated virus pellet was resuspended in phosphate buffered saline (PBS), adding 1/10^th^ of the volume in which the supernatant was mixed with PEG-IT. Virus was then aliquoted and stored at –80°C. The concentration of viral stock was determined by quantifying the amount of ng•p24Gag per mL, using an HIV-1 p24 high sensitivity AlphaLISA Detection kit (Revvity).

### 2.5 p24 Alphalisa Assay

Concentration of p24Gag protein was performed using a HIV-1 p24 High Sensitivity AlphaLISA Detection Kit (Revvity) as done previously^51^. Briefly, all reagents were brought up to RT in low light conditions. A 10X master mix was generated by diluting Anti-HIV-1 p24 Acceptor Beads and Biotinylated Anti-HIV-1 p24 Antibody 1:50 in 1X Alphalisa ImmunoAssay Buffer. Then 5 μL of 10x Master Mix and 5 μL of the supernatant to be assayed were added to each well in a white, half-area 96 well plate (#6002290, Revvity). After addition of samples, human HIV-1 p24 standard was serially diluted according to the manufacturers protocols to generate a standard curve with a range of 0.3 ng/mL to 100,000 ng/mL. Standards were loaded in duplicate, and viral stocks & supernatants were loaded in duplicate at each dilution. After loading, plates were sealed with an opaque, black plate cover (#6005189, Revvity) and incubated for 1 hour at RT on a high-speed shaker (Corning LSE Digital Microplate Shaker, Millipore Sigma) set to 1000 rpm. Following this incubation, SA-Donor beads were diluted 1:100 in 1x AlphaLISA immunoassay buffer and 40 μL of bead dilution was added to each well. Plates were then re-covered with black plate covers and incubated at 1000 rpm for 30 min at RT. Plates were then read at 615 nm on an Enspire Multimode plate reader (Revvity).

### 2.6 Viral Titer Assay

Both parental and Hi-5 GHOST cells were plated at 24 hr prior (Day –1) to starting the viral titer assay. In 6 well plates, GHOST cells were plated at 10^5^ cells/well. In 96 well plates cells, multiple plating densities (0.5 – 11 x 10^3^ cells/well) were tested and a plating density of 3 x 10^3^ cells/well was selected for the majority of studies. In 96 well plates, cells were plated in the center 60 wells of a black walled, transparent bottom 96 well plate (#165305, ThermoFisher). The outer well of each row was filled with Hank’s Balanced Salt Solution (HBSS) (ThermoFisher) to minimize edge effects.

After 24 hrs (Day 0), viral samples to be assayed were prepared in microcentrifuge tubes, diluting the samples in culture medium for either the parental or Hi-5 GHOST cells. Media was then removed from the GHOST cell cultures and replaced with diluted viral samples, adding 1mL per well in 6 well plates and 100 μL per well in 96 well plates. Samples were added to a single well in 6 well plates and in triplicate to 96 well plates. Following inoculation with viral samples, plates were incubated at 37°C and 5% CO_2_ for 2.5 hrs, then media with viral samples was removed and replaced with fresh Complete GHOST culture media, adding 200 μL per well in 96 well plates and 2 mL per well in 6 well plates. Plates were then incubated for 48 hrs at 37°C and 5% CO_2_. After 48 hrs, media was removed, cells were washed once with 100 μL 1x PBS, stained with Hoechst (1:1000 in PBS, ThermoFisher) for 10 min at RT and then washed 2x with 100 μL PBS. After washing, plates were placed in the Cellomics CX7 High Content Screening (CX7) platform (ThermoFisher) and imaged using the 10x objective. Automated imaging acquired 25 fields per well from 96 well plates, with a minimum of 2 wells per sample across the assay. For six well plates, 50 fields were acquired from a single plate per viral stock sample.

Images were analyzed using HCS Studio 2.0 Cell Analysis software (ThermoFisher). The primary measure assessed was the colocalization of blue fluorescence (nuclei) with green fluorescence (infected GHOST cells). Nuclei were identified as primary objects (Region of Interest A Mask Ch 1 (ROIA.Mask.Ch1) using the Hoechst stain in channel 1. Fluorescence due to debris and other auto fluorescent objects were removed by establishing thresholds and segmentation of single nuclei for size and intensity. Objects that were bisected by the edge of the field image were removed to avoid double counting. GFP expression induced by infection of GHOST Hi-5 cells identified secondary objects in channel 2 (ROIA.Target1). Green objects were gated based on intensity, using threshold and segmentation to create masks around the GFP producing cells. Primary objects were used as a guide for object mask creation. The GFP fluorescence threshold for background was defined by GFP fluorescence level in the GHOST cultures inoculated with vehicle. After removing background fluorescence, total nuclei in all fields were enumerated and colocalization analysis was used to define the total number of nuclei within green regions, which were designated as infected cells. Each nucleus was counted as a single cell, enabling determination of the percentage of infected cells. All parameters used in the HCS analysis are described in Table 1.

**Table 1.** High Content Imaging Parameters: Conditions used in HCS software (Cellomics) analysis of images taken by the Cellomics CX7 platform. A Y denotes that this condition was a checkbox.

| Condition | Value |
| --- | --- |
| Ch1: Smoothing (Uniform) | 2-3 |
| Ch1: Thresholding (Fixed) | 1,200.0-2,000.0 |
| Ch1: Segmentation (Intensity) | 150.0-250.0 |
| Object.BorderObject.Ch1 | Y |
| Object.Ch1.Area.Ch1 | 77.55-2432.34 |
| Object.AvgIntensity.Ch1 | 1,400.0-65,535.0 |
| Ch2: Smoothing (Uniform) | 3 |
| Ch2: Thresholding (Fixed) | 1500-2500 |
| Ch2: Segmentation (Intensity) | 6000 |
| Object.Ch2.Area.Ch2 | 483.09-50976.79 |
| Object.Ch2.AvgIntensity.Ch2 | 1531.98-15691.45 |
| ROIA.Mask.Ch1 | CH1-DAPI |

Once HCS analysis was completed, the ROI A Target I Object Count, i.e. the number of cell nuclei colocalized with green fluorescence across all fields in a well, were exported to a spreadsheet in Microsoft Excel (Microsoft, Bellingham, WA) for analysis. This cell-level data contained a binary score for each cell, with a 0 for nuclei not colocalized with green fluorescence and a 1 for nuclei colocalized with green fluorescence. Objects were counted using the countIF function, characterizing an object as an infected cell if it has a colocalization readout value of greater than 0 and if it is part of the total cell population with a score greater than –1. This enabled the inclusion of all 0 values in Excel. Objects were enumerated across all fields of within each well, then combined to generate a single value for the number of infected and total cells in each well. Counts from replicate wells within a single condition were pooled and averaged. Percent infection (equation 1) and subsequently infectious units (equation 2) were then calculated from these data using the following equations:

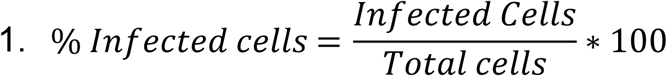

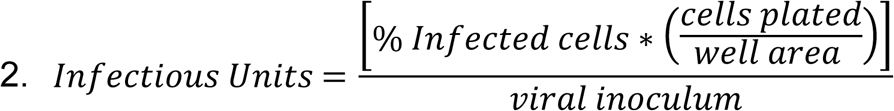

In the second equation, infectious units were defined by cell density per μL. Cell density was defined as the ratio of infected cells to well area (96-well plate is 0.32 cm^2^). For example, if 5 μL of a viral sample were diluted into 100 μL and then used to infect a well containing 3,000 cells and 8% of the cells were infected, you would run the calculation as follows.

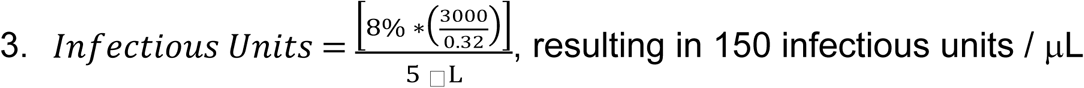

To scale up to infectious units per mL, values are multiplied by a factor of 1000. To calculate the ratio of IU:p24, IU/μL was converted to IU/mL, facilitating the calculation of the IU/24 by comparing like volumes of ng p24 and IU.

### 2.7 Statistics

All data were first evaluated for skewness and normality to determine the distribution of the data. Descriptive statistics (mean, standard deviation (Std. Dev.), standard error of the mean (SEM) and coefficient of variation (CoV)) were defined for each data set. Std. Dev. was used to evaluate viral stocks, while SEM was used to evaluate experiments done using primary human samples. Parametric (t-test, ordinary and repeated measures one-way ANOVA) or non-parametric (Wilcoxon test) tests and correlation analyses (Pearson and Spearman, simple linear regression) were used as appropriate. Outliers were assessed using Grubbs’ and excluded from further analysis. Analyses were set to account for multiple comparisons, and *post-hoc* analyses were performed when appropriate (Tukey’s). All data analysis was performed using GraphPad Prism 9.0 for MacOS (Graphpad, La Jolla, CA). p < 0.05 was considered significant.

## 3. Results

### 3.1 Establishing high content imaging analysis of viral titer

Initial studies evaluated high content analysis of infectious titer using an R5 tropic viral stock, HIV-1_ADA_, generated via infection of CEM•SS cells, titering viral stocks in parental (CD4+/CCR5-) and Hi-5 (CD4+/CCR5+) GHOST cells in 6 well plates. Cells were inoculated with 1, 2.5, 5, 10 or 20μL of HIV-1_ADA,_ stained with Hoescht and analyzed using the CX7 (Figure 2A). Inoculation of Hi-5 cells with vehicle, as well as inoculation of non-CCR5 expressing parental cells, served as controls. Infected Hi-5 cells showed robust GFP expression, while parental cells, which lack HIV-1 co-receptor expression showed no GFP (Figure 2B). Images were analyzed using colocalization of (Hoechst) nuclei (386nm primary channel) and GFP (506nm, secondary channel), showing that the HIV-1_ADA_ stocks contained 351.2-961.2 IU/μL. No statistical differences were observed between the concentrations generated by any of the inoculum volumes used on any of the assay runs performed (Figure 2C, n = 6, rmANOVA; comparison between inoculum volume, F(5,25) =1.131, p = 0.3697; 1μL range 132-718, Std. Dev. 199.2; 2.5μL range 420-951.5, Std. Dev. 212.2; 5μL range 517-778, Std. Dev. 100.2; 10μL range 591-951, Std. Dev. 129.5; 20μL range 618-1370, Std. Dev. 282.9). Notably, this level of variability is at the lower end of the variability seen in comparable TCID_50_ and plaque assays, which generally show 1 to 2 log variation in defined values ^53–55^. The magnitude of the standard deviation was greatest with smaller (1 and 2.5 μL) and larger (20 μL) inoculum volumes than intermediate volumes (5 and 10 μL), so intermediate volumes were used in subsequent experiments.

**Figure 2.**
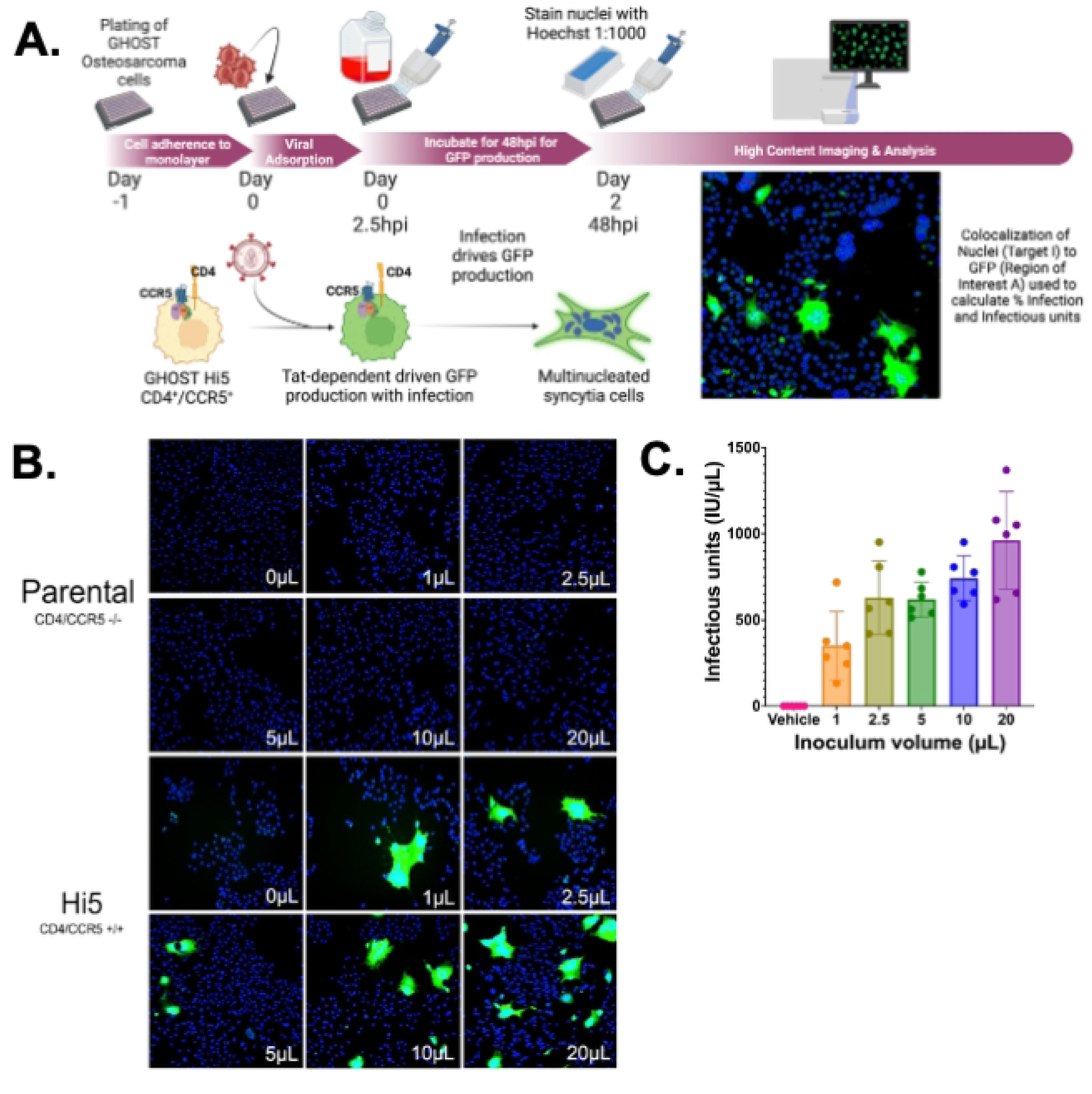
GHOST cells containing a CCR5 receptor can be used to evaluate the presence of infectious virions of HIV-1 from R5-tropic strains. Ghost (3) cells of the genotypes CD4/CCR4^+/+^(Hi5) and CD4/CCR5^-/-^ (Parental) were plated in 6 well plates at 100,000 cells per well and then infected with 0 to 20 μL of HIV-1_ADA_ and incubated for 48hrs at 37°C, as represented in (A.). Cells were washed with PBS and stained with Hoechst (1:1000) and imaged as seen in (B.) at 10x magnification with 50 fields per well. Quantification in (C.) shows that Hi5 cells are infectable, and increase in viral infection with increasing amounts of virus in an n = 6 independent experiments, (range 1μL 132-718, 2.5μL 420-951 5μL, 517-778, 10μL 591-951, 20μL 618-1370, Std. Dev. 1μL 199.2, 2.5μL 212.2 5μL, 100.2, 10μL 129.5, 20μL 282.9, one way ANOVA Treatment, *p-value = 0.0002, F = 21.48, one way ANOVA runs p = 0.3697 F = 1.131).

To adapt the assay to a high content format, Hi-5 cells were plated in black walled, 96-well plates at 500 to 11,000 cells per well. A wide range of cell densities were tested accounting for inconsistencies between plate sizes. Each density was inoculated in triplicate with either 5 μL or 10 μL of HIV-1_ADA_, and plates were imaged at 48 hours post infection. Infection was observed at all cell densities, with minimal background in vehicle wells (Figure 3A). There was a positive, linear correlation between cell density and calculated number of infectious units (Figure 3B, n = 4, Pearson r = 0.9915, Simple linear regression r^2^=0.9831, ****p = 3.421×10^−10^). Data from wells with 500 and 1,000 cells per well were less consistent, potentially due to sparseness that generated inconsistent infection between wells within a single inoculation condition. Densities greater than 5,000 cells per well were also inconsistent, likely due to the formation of clusters that were difficult to discriminate as individual cells. The number of infectious units (IU/μL) did not differ between wells with 2,000 to 5,000 cells per well (Supplemental Table 1, One-way ANOVA ****p = <0.0001, F (11, 36) = 17.06, Tukey’s *post-hoc*). The lowest percentage variation in IU/μL resulted from data with 2,000 or 3,000 cells per well (Supplemental Table 2), so a density of 3,000 cells per well was selected for further assays in 96 well plates.

**Figure 3.**
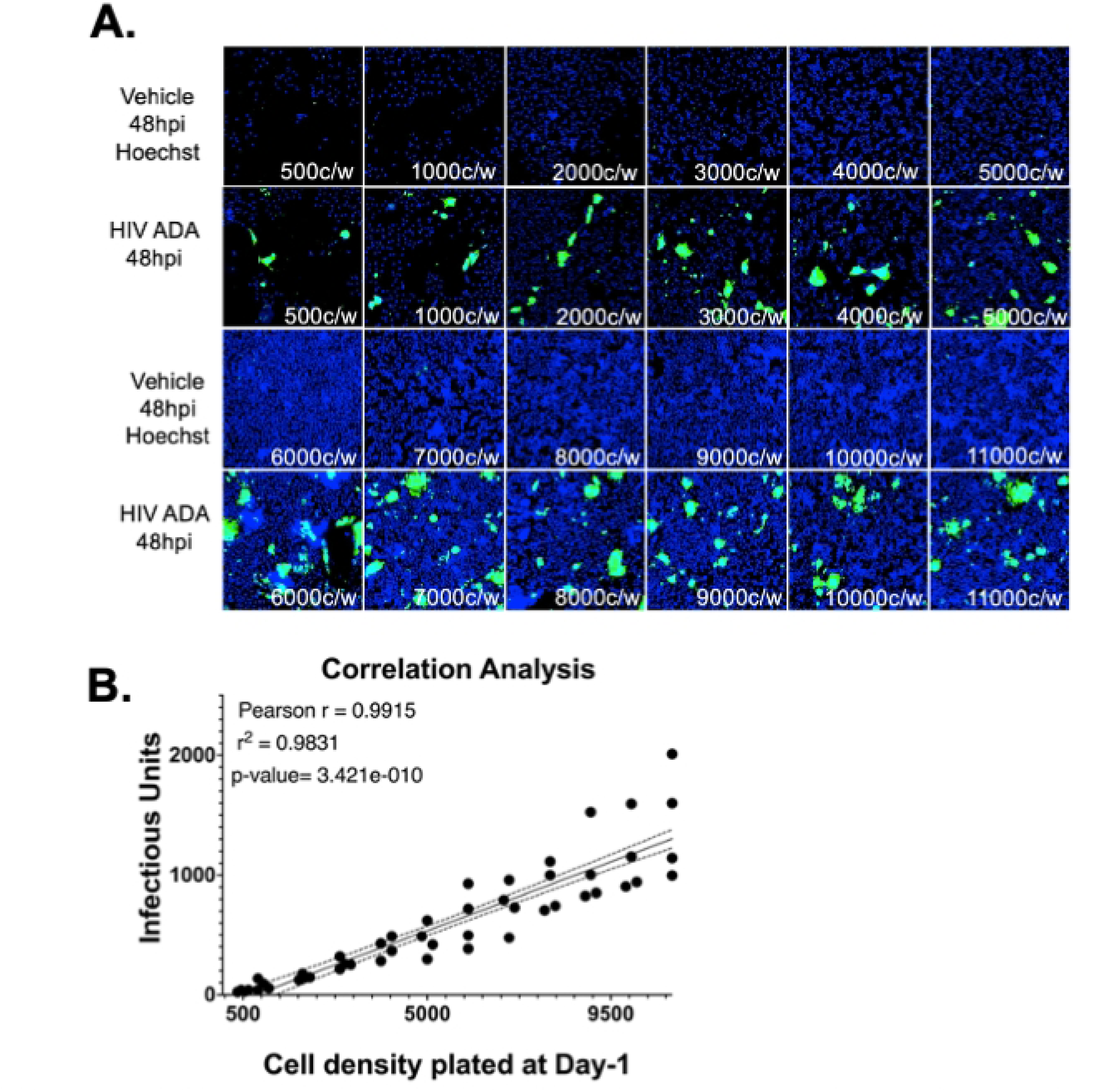
The viral titer/ GHOST cell assay can be optimized and used in a high throughput manner in 96-well plates based on initial cell density. (A.) Hi-5 cells were plated at 500 to 11,000 cells/well and infected in triplicate with 5 μL or 10 μL of HIV-1_ADA_. Infection was observed at all densities, with minimal signal in vehicle controls. (B.) A strong linear correlation was found between cell density and infectious units (n = 4, Pearson r = 0.9915, R² = 0.9831, ****p = 3.421 × 10⁻¹⁰). Variability was highest at the lowest (≤1,000 cells/well) and highest (≥5,000 cells/well) densities. Infectious titers did not differ significantly between 2,000 to 5,000 cells/well (One-way ANOVA, ****p < 0.0001, F(11,36) = 17.06; Tukey’s post hoc).

### 3.2 Defining the assay window

To define an optimal assay window, HIV-1_ADA_ stocks were diluted to concentrations between 0.1 ng•p24/mL to 100 ng•p24/mL in complete GHOST Medium and then assessed for IU/μL. Three replicate wells were inoculated with each concentration, repeating the assay in 5 independent experiments and then pooling the mean IU/μL for each concentration across all 5 assays. Infection was observed in response to inoculation with all HIV-1_ADA_ concentrations (Figure 4A-B, One-way ANOVA, ****p = <0.0001, individual p-values determined by Tukey’s multiple comparisons) and the IU/μL was significantly correlated concentration of ng•p24/mL (Figure 4C, Pearson Correlation, r = 0.9308, **** p = 0.00003192, r^2^ = 0.8664). Despite detectable infection, the lowest inoculum concentrations (0.1 ng•p24/mL, 0.25 ng•p24/mL and 0.5 ng•p24/mL) did not induce fluorescence in all 5 infectious titer assays, as shown by the points below the dashed line in Figure 3B. Lack of detectable infection in response to very low inoculum is an issue present in all reporter assays, and we found that this can be accommodated by increasing the probability of detecting infectivity by testing inoculum with very low p24 concentrations in a larger number of wells.

**Figure 4.**
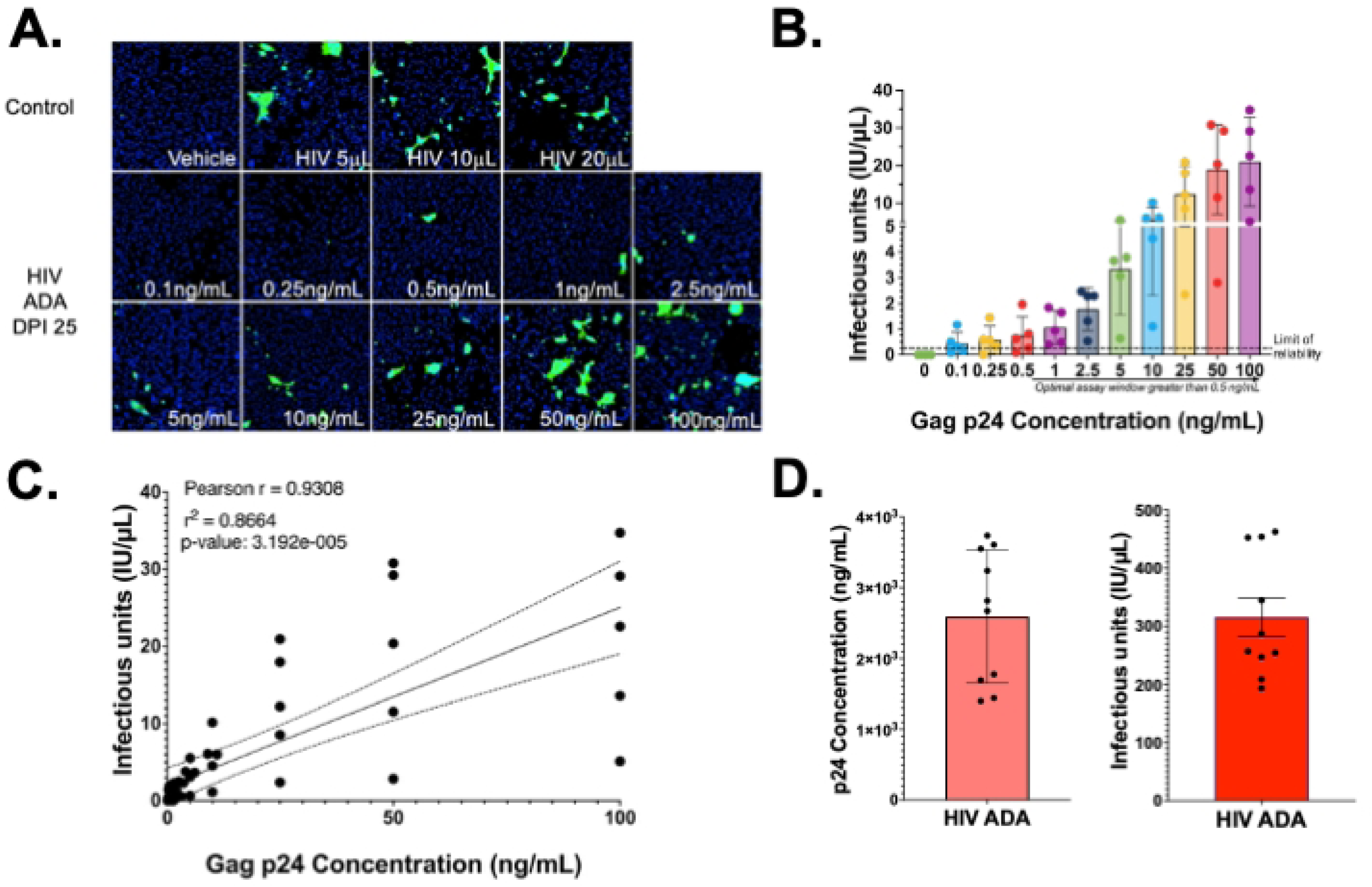
The viral titer assay window of reliability is comparable to measurement of p24. (A.) Representative images (B.) quantification of an n=5 independent experiments of infecting Hi5 cells with 0-100ng/mL p24 concentration of HIV-1_ADA_ showed that as p24 increases infectious unit production also significantly increases with a limit of reliability (dashed line) at 0.5ng/mL (One-way ANOVA p-value <0.0001, Tukey’s multiple comparisons p-values depicted <0.001, for other p-values see Supplemental Table 3). (C.) Pearson correlations of infectious units and p24 showed a strong positive linear correlation between the two variables (Pearson r = 0.9308, p-value 3.192 × 10^−5^, r^2^ = 0.8664). (D) Quantification of biological replicates from two HIV-1_ADA_ stocks using viral titer (315.9 IU/μL, Std. Dev. = 105.1, CoV = 33.26%) and p24 alphaLISA (2.593 × 10³ ng p24/mL Std. Dev. = 939.3, CoV = 36.23%).

**Figure 5.**
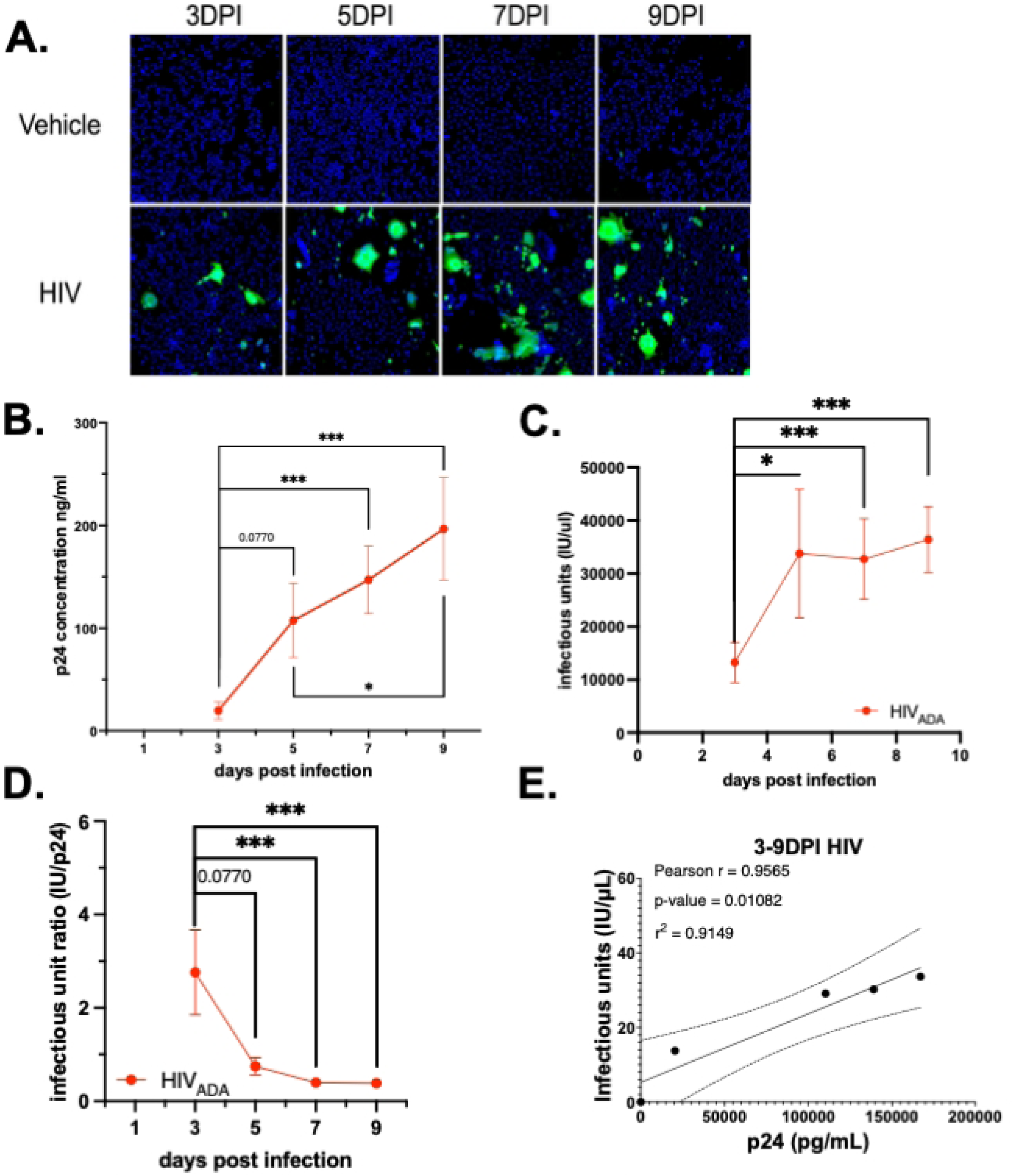
The viral titer assay can evaluate infectious virion production in hMDMs. (A) Representative images of viral titer assay from cell treated with supernatants from HIV-1_ADA_ – infected hMDM (1 ng p24/mL) at 3, 5, 7, and 9 dpi. (B) analysis of p24 secretion (C) infectious titer (D) IU:p24 ratio

To evaluate assay consistency, the IU/μL of a single HIV-1_ADA_ viral stock was assessed across 10 independent assays. The HIV-1_ADA_ stock has an infectious titer of 315.9 IU/μL (Fig. 4D, n=10, Std. Dev. 1.051 x 10^2^ IU/μL, CoV 33.26%). This value was compared with the ng•p24/mL of that stock quantified p24 AlphaLISA, which was determined to be 2.593 x 10^3^ ng•p24/mL (Fig. 4D, n=10, Std. Dev. 9.3929 x 10^2^ ng•p24/mL, CoV 36.23%). The p24 AlphaLISAs and high-content viral titer assays had very similar standard deviation and coefficients of variation, indicating that the two assays have similar consistency, although detection of p24 secretion by AlphaLISA may be more sensitive in samples with low concentrations of virus. Both assays evaluate the same supernatants but examine distinct stages of the viral replication cycle, thus running them in parallel enables assessment of changes in capsid production and virion maturation in the same samples.

### 3.3 Evaluating changes in macrophage infection using infectious titer

Prior research in the Gaskill laboratory has assessed HIV-1 infection in hMDM via evaluation of viral entry or measurement of secreted p24 by AlphaLISA^51, 56^. To demonstrate the utility of the infectious titer assay within this existing experimental pipeline, recent and archival supernatants collected from HIV-1_ADA_ (1 ng•p24/mL)-infected primary human monocyte derived macrophages (hMDM) obtained from 13 donors at 3, 5, 7 and 9-days post-infection (dpi) were quantified for IU/μL (Figure 5A). This was then compared with prior analysis of p24 secretion in these samples. The pooled p24 secretion in these samples showed a consistent increase in p24 secretion over time (Figure 5B, n=14, Mean of Day 3 19.46 ng/mL, SEM 8.418 ng/mL, CoV 161.8%; Day 5 Mean 125.1 ng/mL, SEM 35.99 ng/mL, CoV, 126.9%; Day 7 Mean 147.1 ng/mL, SEM 32.73 ng/mL, CoV 83.23%; Day 9 Mean 167115ng/mL, SEM 49.71 ng/mL, CoV 94.6%, rm one-way ANOVA, Friedman test *p<0.05, days 3-5 p=0.0770, days 3-7 ****p<0.0001, days 3-9 ****p<0.0001). The high levels of variability in these infections are expected in infection of primary human MDM^57^.

Assessment of infectious titer in these same supernatants also showing a consistent increase of IU/μL over time (Figure 5C, n=14, Day 3 Mean 13267 IU/μL, SEM 3828 IU/μL, CoV 107.9%; Day 5 Mean 33787 IU/μL, SEM 12124 IU/μL, CoV 134.3%; Day 7 Mean 28360 IU/μL, SEM 7580 IU/μL, CoV 86.67%; Day 9 Mean 36.396 IU/μL, SEM 6204 IU/μL, CoV 63.78%, rm one-way ANOVA, Friedman test *p<0.05, days 3-5 **p=0.0077, days 3-7 **p=0.0015, days 3-9 ****p<0.0001). The ratio of IU to p24, which represents the number of infectious virions per unit of p24Gag protein, showed that the highest ratio IU:p24 occurs early in infection (Figure 5D, n=14, Day 3 Mean IU:p24 2.762, SEM 0.909, CoV 123.2%, Day 5 mean IU:p24 0.748, SEM 0.1835, CoV 91.84%, Day 7 Mean IU:p24 0.397, SEM 0.0904, CoV 85.25%, Day 9 Mean IU:p24 0.339, SEM 0.091, CoV 88.84%; Friedman test *p<0.05, days 3-5 p=0.0770, days 3-7, ***p=0.0001, days 3-9 ****p<0.0001, days 5-7, p=0.942, days 5-9, p=0.9350, days 7-9, p>0.9999, r^2^=0.3406.

Notably, the IU:p24 ratio was significantly reduced by 9 dpi, indicating that more p24 was associated with each infectious virion at that timepoint. This suggests that the efficiency of virion production decreases as infection progresses. Correlating the average p24/mL and IU/μL at days 3, 5, 7 and 9 showed a linear correlation between these factors (Figure 5E; HIV-1, Pearson r = 0.9565, *p = 0.0108, r^2^= 0.9149). This confirmed that, at least in hMDM, evaluation of viral titer reflects the same trends as measurement of p24 secretion. The relatively smaller coefficients of variation in IU/μL relative to ng•p24/mL suggest that the viral titer assay may be particularly useful when evaluating infection in data sets with large variability, such as infection of primary cells from multiple donors. Further, combining analysis of p24 secretion and infectious titer assays within the same experimental pipeline has the potential to provide important nuance to the understanding of viral replication kinetics, viral capsid secretion and virion maturation.

### 3.4 Assessing viral infectious across distinct strains of HIV-1

To demonstrate that the assessment of viral titer is consistent across distinct preparations and strains of HIV-1, the infectious titer assay was used to analyze viral stock generated from multiple strains of R5 tropic HIV-1 (Figure 6A). The first strain tested was HIV-1_YU2_, a laboratory adapted R5-tropic strain of HIV-1 isolated from brain tissue. For this analysis, viral stocks were generated from two distinct preparations of HIV-1_YU2_, designated HIV-1_YU2**A**_ and HIV-1_YU2**B**_ (Figure 6B). While the clones used to generate HIV-1_YU2**A**_ and HIV-1_YU2**B**_ are genetically identical (Supplemental Figure 2), the plasmids containing the each YU2 clone were obtained from two different sources. The HIV-1_YU2**A**_ was obtained from then NIH HIV-1 Reagent program and amplified by Genescript, while the HIV-1_YU2**B**_ plasmid was a generous gift from the Bar lab at the University of Pennsylvania. Viral stocks were generated from each plasmid using Lenti-X cells and evaluated for p24 concentration (ng•p24/mL) (Figure 6C, HIV-1_YU2**A**_ (n=3) Mean=10689 ng/mL, Std Dev. 13736, CoV 128.5%; HIV-1_YU2**B**_ (n=4) Mean=7923 ng/mL, Std. Dev. 12442, CoV 157%, unpaired t-test, p=0.7914), and infectious units (IU/μL) (Figure 6D, HIV-1_YU2**A**_ (n=3) Mean=69.5, Std. Dev 78.14, CoV 112.4%; HIV-1_YU2**B**_ (n=4) Mean = 131.5, Std. Deviation 100.5, CoV 76.40%, unpaired t-test, p=0.59). To determine the efficiency of infectious virus production, i.e. the amount of p24 secreted per infectious unit, the IU/μL was divided by the amount of ng•p24/mL to generate a ratio of IU:p24 (Figure 6E, HIV-1_YU2**A**_ (n=3) Mean = 0.0025, Std. Dev. 0.003, CoV 111.9%; HIV-1_YU2**B**_ (n=4) Mean = 0.095, Std. Dev. 0.1654, CoV 173.4%, unpaired t-test, p=0.52) for both viral stock preparations. While the two preparations had comparable ng•p24/mL concentrations, the stock derived from HIV-1_YU2**B**_ showed a greater amount of IU/μL and a slightly higher IU:p24 ratio, which could indicate a more infectious stock.

**Figure 6.**
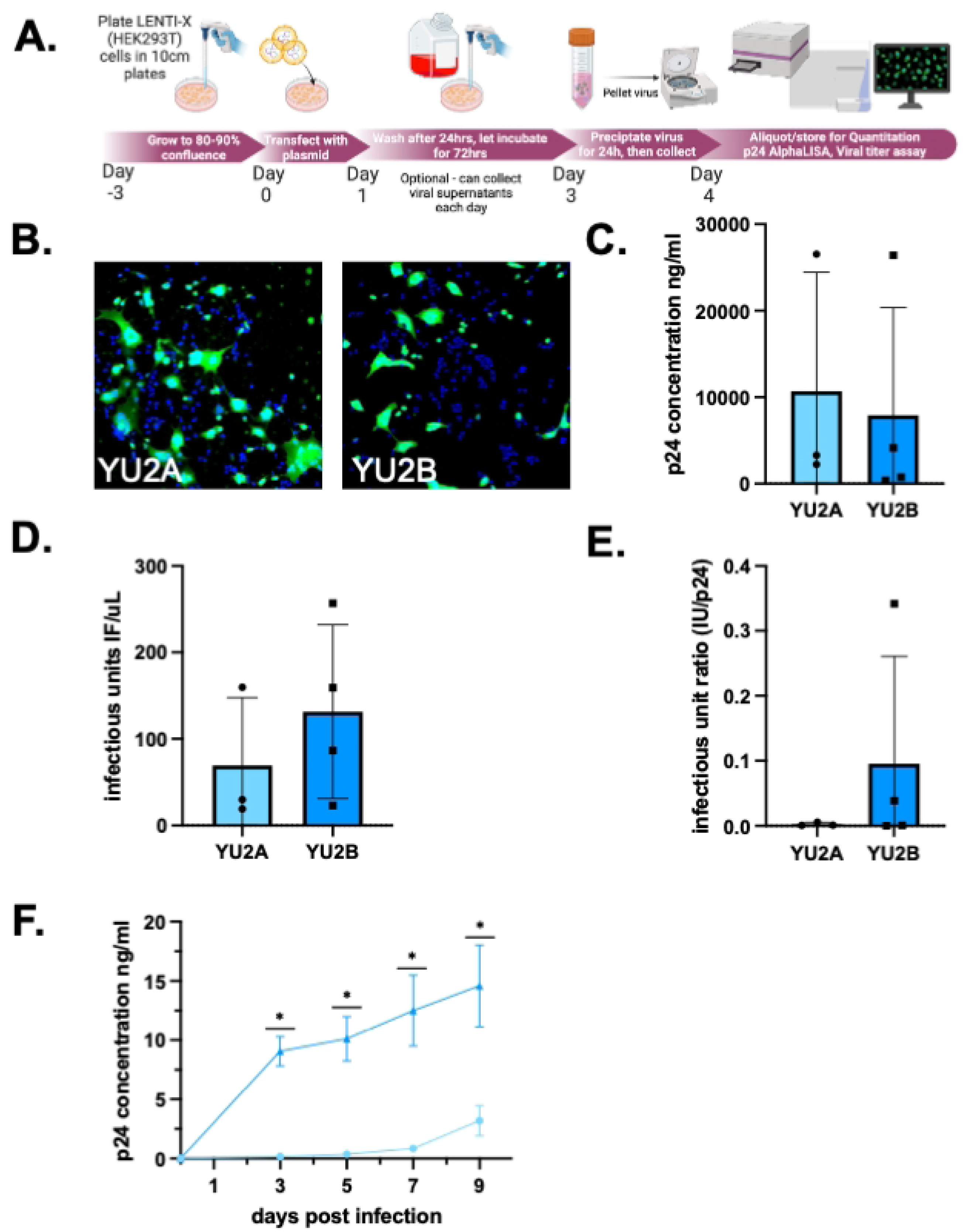
Infectious unit ratio highlights differences between viral stocks of the same strain. (A) Schematic overview of lentiviral transfection to generate to R5-tropic strains of HIV-1. (B) Representative images of viral titer assay from two stocks of HIV-1_YU2_ that are genetically identical but obtained from distinct sources. (C) Quantification of p24 levels (ng/mL) (D) Measurement of infectious titer (IU/μL) in YU2B compared to YU2A (E) The IU:p24 ratio was calculated to assess the relative efficiency of infectious virus production (F) hMDMs from matched donors were infected with equivalent p24 concentrations (20 ng/mL) from YU2A or YU2B stocks. Longitudinal measurement of p24 secretion.

This suggests that the viral preparation using HIV-1_YU2**B**_ resulted in a more efficient production of infectious virions, as a smaller amount of p24 was associated with each infectious unit. Similarly, it suggests that the viral stock generated using HIV-1_YU2**A**_ resulted in the production of a higher number of defective, i.e. non-infectious, virions as a greater amount of p24 was associated with each infectious unit. To confirm this, HIV-1_YU2**A**_ and HIV-1_YU2**B**_ were used to infect hMDM from the same donors, inoculating with identical amounts of p24 (20ng/mL) from each viral stock. Longitudinal analysis of these infections measuring both secreted p24 and IU/μL showed that HIV-1_YU2**B,**_ the stock with the higher IU:p24 showed more robust infection in primary hMDMs, as predicted by the IU/μL and IU:p24 ratio (Figure 6F, HIV-1_YU2**A**_ (n=6) DPI3 Mean = 148.3 ng•p24/mL, SEM 81.84, CoV 146%, DPI5 Mean = 354.9 ng•p24/mL, SEM 151.5, CoV 104.6%, DPI7 Mean = 853.2 ng•p24/mL, SEM 306.9, CoV 88.21%, DPI9 Mean = 3188.0 ng•p24/mL, SEM 1272, CoV 89.21%; HIV-1_YU2**B**_ (n=7) DPI3 Mean = 9065 ng•p24/mL, SEM 1243, CoV 36.29%, DPI5 Mean = 10110 ng•p24/mL, SEM 1882, CoV 45.57%, DPI7 Mean = 12510 ng•p24/mL, SEM 2997, CoV 63.39%, DPI9 Mean = 14590 ng•p24/mL, SEM 3441, CoV 57.76%, mc one-way ANOVA *p<0.05, DPI3 **p=0.0041, DPI5 **p=0.0036, DPI7 ***p=0.003, DPI9 **p=0.0011; HIV-1_YU2**A**_ slope = 286.2, HIV-1_YU2B_ slope = 1541, nonlinear fit ***p=0.0004). In addition to showing a statistically significant difference in the values of ng•p24/mL at each time point, the HIV_YU2B_ stock also showed an increase in the rate of infectivity as measured by the slope of the line. This analysis highlights the value in using multiple outputs to characterize viral stocks – as the production of “non-infectious” viral particles is common in HIV-1 – and shows the capacity of the combined assays to increase both experimental efficiency and our understanding of HIV-1 kinetics.

To further test this assay, assessment of ng•p24/mL and IU/μL was performed on viral stocks generated viruses obtained from three distinct participants (Figure 7A). The viruses used include the transmitted founder clinical isolate HIV-1 THRO (Genbank: JN944930.1^58^), the rebound clinical isolate 9201R1 (Genbank: MT929391^59^) and a chronic clinical isolate CH167 (Genbank: KC156213, Hora et al. Unpublished). All stocks showed detectable levels of p24Gag (ng•p24/mL) (Figure 7B n=2 CH167 Mean = 14172.413 ng/mL, Std. Dev. 17906.206, CoV 236.346%; n=4 9201R1 Mean = 12086.109 ng/mL, Std. Dev. 15124.893, CoV 125.3%; n=4 THRO Mean= 1828.571 ng/mL, Std. Dev. 2082.55, CoV 113.889%) and IU (Figure 7C, n=2-3; CH167 Mean = 63.956 I IU/μL, Std. Dev. 51.98, CoV 81.28%; 9201R1 Mean = 118.233 IU/μL, Std. Dev. 56.02, CoV 47.378%; THRO Mean= 189.9 IU/μL, Std. Dev. 211.62, CoV 111.44%). Notably, although CH167 did have quantifiable p24 and IU, it had the lowest IU count of the 3 participant-derived viruses.

**Figure 7.**
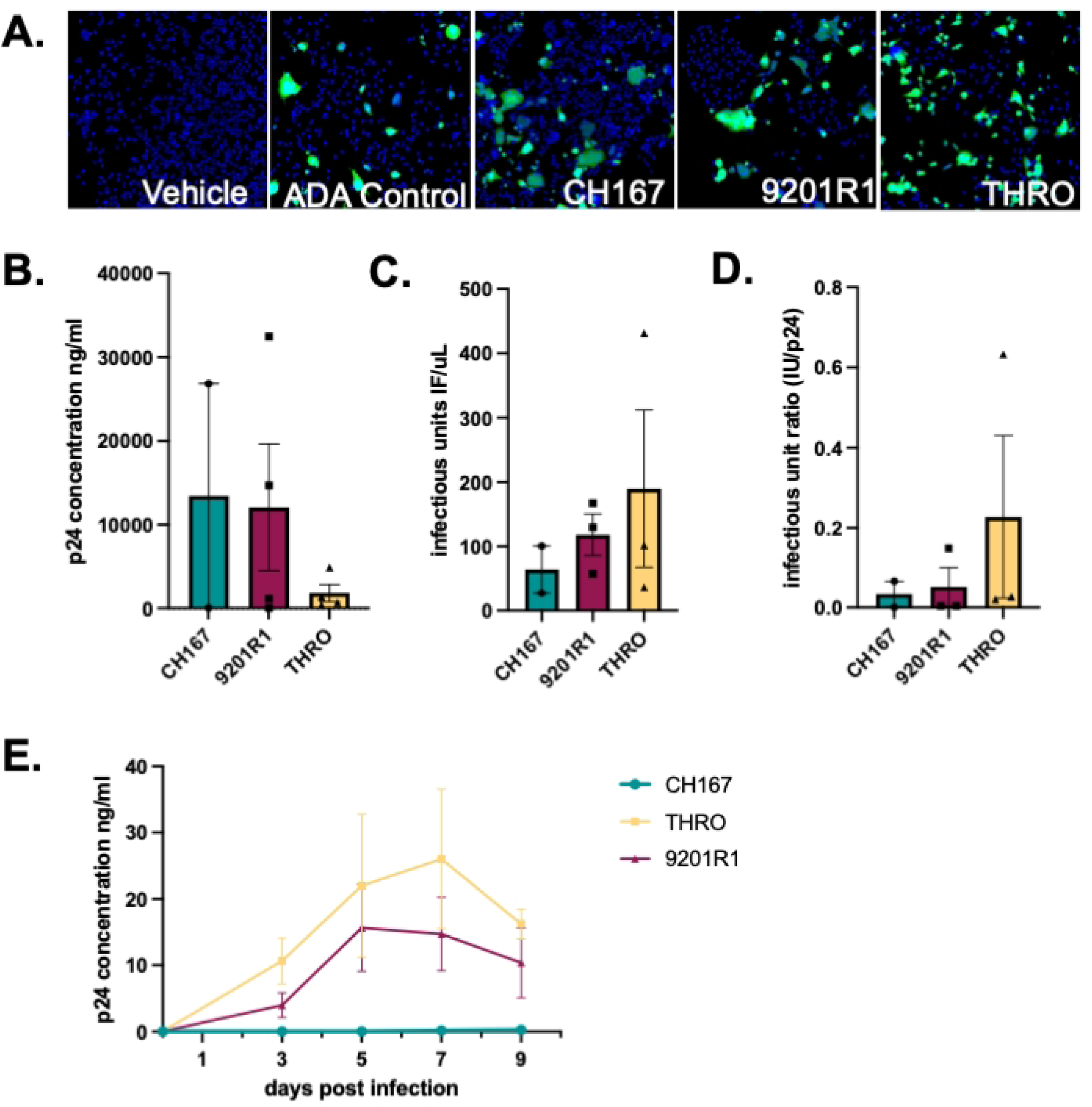
Characterizing viral output from infection of primary human monocyte derived macrophages. (A.) Representative images from viral titer assay of participant-derived viral stocks generated through lenti-viral transfections (B) Quantification of p24 (ng/mL) in viral supernatants demonstrated detectable levels in all stocks (C) Infectious titer (IU/μL) varied between stocks (D) Calculation of the IU:p24 (E) hMDMs were infected with equal p24 input from each viral stock (20 ng/mL), and infection was assessed longitudinally by p24 secretion.

Although all stocks had measurable levels of p24 and IU, there was substantial variability between strains, with a very low IU:p24 ratio for CH167, a higher but still relatively low ratio for the 9201R1 stocks, and the highest IU:p24 ratio for the THRO virus (Figure 7D, n=2-3; CH167 Mean = 0.034, Std. Dev. 0.046, CoV 137.2%, 9201R1 Mean = 0.05, Std. Dev. 0.08, CoV 160.1%, THRO Mean = 0.227, Std. Dev. 0.35, CoV 154.98%).

Analysis of hMDM inoculated with all three viral stocks reflected pattern, as 9201R1 and THRO appear to be more infectious than CH167 in regard to longitudinal production of p24 as a measure of infection (Figure 7E; CH167, n=1, DPI3 Mean = 0.01195 p24•ng/mL, DPI5 Mean = 0.0245 p24•ng/mL, DPI7 Mean = 0.1367 p24•ng/mL, DPI9 Mean = 0.2508 p24•ng/mL; 9201R1 n=4, DPI3 Mean = 3.968 p24•ng/mL, SEM 1.877, CoV 94.62%, DPI5 Mean = 15.64 p24•ng/mL, SEM 13.08, CoV 83.64%, DPI7 Mean = 14.73 p24•ng/mL, SEM 5.548, CoV 75.32%, DPI9 Mean = 10.39 p24•ng/mL, SEM 5.276, CoV 101.6%; THRO n=4, DPI3 Mean = 10.66 p24•ng/mL, SEM 3.485, CoV 65.35%, DPI5 Mean = 21.97 p24•ng/mL, SEM 10.78, CoV 98.1%, DPI7 Mean = 26.02 p24•ng/mL, SEM 10.51, CoV 80.79%, DPI9 Mean = 16.22 p24•ng/mL, SEM 2.239, CoV 27.61%). As noted above, CH167 had the lowest IU:p24 ratio, and IU count overall – and did not show detectable infection. These data highlight the variability associated with viral stock preparation and demonstrate the utility of the high throughput adaptation of the infectious titer assay. These data also showcase the capacity to combine these assays to generate an IU:p24 ratio that may more wholistically reflect the ability of viral stock to infect primary cells with HIV-1.

## 4. Discussion

Quantification of HIV-1 infection and spread in cell culture is critical to understanding HIV-1 replication dynamics. The assay developed in this study adds to the existing toolkit in this area, adapting a pre-existing reporter assay using GHOST cells, to a high-content, high-throughput pipeline, enabling more rapid, direct measurement of infectivity in large numbers of samples while minimizing experimenter bias. While these studies utilize GHOST cells, many common reporter lines, such as TZM-bL or CEM-GFP^60^ could be adapted to this protocol as long as the growth and analysis are properly optimized and used on a high-throughput instrument. This study specifically assessed R5 strains of HIV-1 and therefore used Hi-5 GHOST cells, which express the CCR5 co-receptor.

Although widely used, reporter assays are not often engaged to evaluate large numbers of samples across time or many different conditions. This type of analysis is particularly important when evaluating longitudinal changes in viral replication in response to different stimuli, such as substances of misuse or ART. A major hurdle for these studies is that evaluation of infectivity directly via plaque assays, TCID_50_ or reporter cell lines^14, 33, 36^ is often time and labor intensive and risks experiment bias^18, 61, 62^. To address this, we adapted the use of the GHOST reporter cells to the CellInsight CX7 High Content Analysis Platform. This automated, high-throughput confocal microscopy imaging system enables rapid, unbiased, automated focus and image acquisition in multiple channels enabling quantification a number of cell-based parameters (cell size, number, and intensity of stain in different channels, etc..)^63–68^. Using the CX7 to assess GFP allows for direct visualization and quantification of infection in existing supernatants. This approach allows direct comparison of p24 and viral titer, enabling a deeper interrogation of the relationship between IU and p24 measurements in subsequent analyses.

Comparison of IU/μL with the p24•Gag/mL in defined dilutions of HIV-1_ADA_ stock demonstrated a statistically significant correlation between amount of p24 and infectivity of our HIV-1_ADA_ stock. This observation was confirmed by a strong correlation between p24 secretion and IU/μL counts in supernatants isolated from infected hMDM. This indicates that for HIV-1_ADA_, measurement of p24 concentration is a reasonable surrogate, although not a complete substitute for, measurement of infectivity. Notably, prior comparison of p24 ELISAs with TCID_50_ assays of direct infectivity did not show this correlation^14^, which is surprising given the common use of secreted p24 as a surrogate for HIV-1 infection *in vitro* ^11, 12, 15, 69–75^. The reason for the lack of correlation in prior assays is not clear. It may be due to the smaller dynamic range in prior assays^14, 73, 75^, the presence of supernatant factors, such as biotin in RPMI, that could also interfere with capture-antibody binding in specific ELISAs, or issues associated with cell density that hindered accurate detection of infection and cell-to-cell spread in different reporter lines. The utility of determining the IU:p24 ratio is that neither separate evaluation of viral capsid secretion (p24 AlphaLISA) or mature virions (viral titer assay) provide a full picture of viral kinetics. The IU:p24 ratio in HIV-1-infected hMDMs (Fig. 5D) shows that each infectious unit is associated with less p24 early in after inoculation, i.e. a high IU:p24 ratio. This ratio decreases over time, indicating that the amount of p24 secreted for each infectious virion is increasing. Our combined analysis of p24 secretion and viral titer support this, and very interestingly suggest that the amount of virus-like particles (VLPs) increases over time but the amount of infectious virions does not stay consistent with this production. This is likely because HIV-1 replication produces many non-infectious VLPs, due to the lack of viral genomic material or properly spliced or functional viral proteins^76^.

Other studies confirm that only a small number of secreted virions are infectious (around 1% of all virions), and importantly, the proportion of non-infectious virions may not be uniform across all infection conditions or viral variants ^70, 71, 74, 77, 78^.

This highlights the inefficiency in the production of infectious HIV-1 virions. It also suggests that evaluation of p24 alone may result in an under-or over-estimation of the amount of mature, infectious virions produced in a particular culture condition. Further, using p24 as a surrogate for viral replication and infectious capacity may be an incomplete measurement, as shown in Figure 6 comparing the two independent preparations of HIV-1_YU2_, HIV-1_YU2**A**_ and HIV-1_YU2**B**_. These data show that the amount of p24 in the viral stock does not correlate with infectivity in hMDM. Characterizing these stocks by p24 concentration alone would suggest that they have similar infectivity. However, by using measurement of both p24 secretion (ng/mL) and infectious titer (IU/μL), we can define the infectious ratio (IU:p24). This showed that HIV-1_YU2**B**_ had a slightly higher IU count and a slightly lower p24 concentration but a much higher IU:p24 ratio, suggesting it was more infectious. Indeed, inoculation of hMDM with both HIV-1_YU2**A**_ and HIV-1_YU2**B**_ showed that HIV-1_YU2**B**_ was much more infectious than HIV-1_YU2**A**_, as it induced an accelerated infection of higher magnitude.

These data suggest that it is important to define the infectivity of a viral stock as precisely as possible. This is particularly important during infection of primary human immune cells such as T-cells and hMDM, as variations in stock infectivity may be compounded by the high variability in infection in primary human cultures ^56, 57, 72, 79–89^. This variability adds a substantial limitation to studying HIV-1 infection in primary human cells, and highlights the value of assessing viral dynamics at multiple points in the HIV-1 life cycle to develop a more consistent, nuanced and holistic view of infection *in vitro* and *ex vivo* ^57, 77, 79, 84, 88^. As an example, the analysis of viral titer in hMDM supernatants that had already been assessed for p24 concentration (Figure 5) provided additional nuance to *in vitro* replication studies. These data showed that the production of infectious virions paralleled the trends seen for secretion of viral protein over time in these experiments, confirming that the secreted p24 represented production of infectious virus. This also showed that the p24 AlphaLISA being used is a good proxy for infectivity in infections of MDM with HIV-1_ADA_. This parallel may differ in distinct conditions or time points, and combining the assessment infectivity per amount of viral protein could providing a more consistent baseline measurement of infection across multiple donors. This measurement could also be very useful evaluating treatments that disrupt later stages of the viral lifecycle. This includes antiretrovirals like capsid or protease inhibitors, as well as stimuli that may disrupt viral maturation. Further, our analysis also showed that p24 (ng/mL) and infectious titer (IU/μL) are not necessarily consistent across stock preparation methods, further highlighting the importance of thoroughly evaluating infectivity.

Running both assays in parallel on the same supernatant samples provides a powerful means to evaluate discrete steps of the HIV-1 replication cycle—specifically, capsid production and virion maturation. This dual approach could be particularly informative when distinguishing between virus particle release and maturation. For instance, unchanged or elevated p24 levels but significantly reduced infectivity could indicate that virion release is intact, but that the maturation process was compromised. Conversely, decreases in both p24 and infectivity could suggest impairments at earlier stages such as assembly or budding. Assessing both aspects of the replication cycle within the same biological samples controls for inter-sample variability and enables a more precise comparison of how specific interventions affect the sequential stages of virion release and virus maturation. This strategy is particularly relevant in the context of therapeutic development. Several classes of antiretrovirals, including maturation inhibitors and capsid-targeting compounds, exert their effects downstream of particle release. A single assay focused solely on p24 levels would fail to detect these defects in maturation, again showing the utility of incorporating both assays into the analysis of viral fitness and replication.

One possible limitation to this assay is that it may be less sensitive to low levels (< 0.5 ng/mL) of viral replication than p24 AlphaLISAs, as these levels were not consistently detected across replicate wells (Figure 4). Alternatively, this might suggest that at lower p24 levels, the majority of these p24 units are not associated with infectious virions. This limitation could reduce the utility of these tests when evaluating the amount of virus in ART treated cultures, or when using the assay to evaluate outgrowth of virus from patient derived samples that produce lower levels of HIV-1, as in a quantitative viral outgrowth assays (QVOA)^90^. This concern can be addressed by increasing the number of infected wells to better capture infrequent infections, as well as by using viral titer assay in parallel with evaluation of p24 release. It may also be possible to increase the uptake of viral particles by utilizing polybrene which has been shown to enhance the efficiency of transduction of viral particles^91^.

These data show the development of a consistent, high-throughput, assay that integrates well with existing assays used to measure HIV-1 infection and can be easily modified to use multiple types of indicator lines. These data indicate that evaluation of viral titer in parallel with analysis of p24 secretion or staining of infected cells could improve evaluation of changes in infection dynamics not examined by looking at p24 concentration alone, i.e., measurement of efficiency of the egress and maturation of released particles into infectious virions ^92–95^. In summary, the high content viral titer assay can consistently and rapidly screen of large numbers of samples and adds a useful tool to the already expansive HIV-1 toolkit for the assessment of viral infectivity and the evaluation of viral replication dynamics. The data presented using this assay underscores the utility of parallel assay design to capture aspects of HIV-1 replication and highlight the importance of addressing specific lifecycle stages when attempting to better understand viral kinetics.

## Acknowledgements

We would like to thank Drs. Zachary Klase, Rebecca Veenhuis, and Alison Carey, as well as all the members of the Gaskill and Nonnemacher labs, for extensive discussion, advice and consideration of the material that substantively helped to shape this paper.

## Funding Sources

This work was supported by grants from the National Institutes of Drug Abuse (R01DA057337, R61DA058501 to PJG), the National Institutes on Mental Health (K01MH132466 to SMM, T32MH079785 fellowship to ENB) and the National Institute of Neurological Disorders and Stroke (R01NS089435 to MRN). Additional funding from the Departments of Microbiology and Immunology and Pharmacology and Physiology at the Drexel University College of Medicine.

## Notes

### Competing Interest Statement

The authors have declared no competing interest.

